# An annotated chromosome-level genome assembly of the Adzuki bean borer *Ostrinia scapulalis* (Lepidoptera: Crambidae)

**DOI:** 10.64898/2025.12.26.696569

**Authors:** Pierre Nouhaud, Sabine Nidelet, Fabrice Legeai, Philippe Audiot, Joanna Lledo, Sophie Valière, Carole Iampietro, Guy Perez, Nawelle Malinge, Camille Meslin, Emmanuelle Jacquin-Joly, Vincent Calcagno, Réjane Streiff

## Abstract

The genus *Ostrinia* (Lepidoptera: Crambidae) comprises two of the most important maize pests worldwide, the Asian and the European corn borers. Here, we present an annotated, chromosome-level genome assembly for the most closely related species, the Adzuki bean borer *Ostrinia scapulalis*, which feeds on various dicotyledon plants. The previous reference genome assembly for this species was generated from short-read sequencing data, resulting in high fragmentation and low completeness. Combining PacBio long read and Hi-C sequencing, we generated a 476 Mb genome with 48 contigs organised into 31 chromosomes (30 autosomes and one Z sex chromosome), with a contig N50 of 16.1 Mbp and BUSCO completeness exceeding 98%. We further combined published and novel RNA-seq data encompassing multiple life stages, tissues, and sexes to annotate 14,261 gene models, reaching a proteome BUSCO completeness of 95%. This highly contiguous reference genome assembly provides a much-improved resource for carrying comparative genomic approaches and better understanding speciation and host plant adaptation in the *Ostrinia* genus.

## Introduction

Phytophagous insects are outstanding systems for studying adaptation and diversification. First, their intimate associations with host plants expose them to strong and predictable ecological pressures, such as plant defensive chemistry, nutritional heterogeneity, and phenological variation (Futuyma and Moreno 1988; Futuyma and Agrawal 2009; Forister et al. 2012). Second, host shift events, which are frequent in this group (Jaenike 1990; Forbes et al. 2017), often require substantial physiological and behavioral changes. These create opportunities for rapid ecological divergence and the formation of host-associated populations, which could lead to the emergence of reproductively isolated units (i.e., speciation events, Drès and Mallet 2002). Third, usually combining short generation times and large effective population sizes, phytophagous insects may also respond quickly to selection (McCulloch and Waters 2023). Together, these features make phytophagous insects powerful models for investigating the genetic architecture of adaptation (Simon et al. 2015), the mechanisms underlying speciation (Jousselin and Elias 2019), and the evolutionary forces that shape biodiversity (e.g., Breeschoten et al. 2022). In addition to their value as model organisms, many phytophagous insects are major agricultural and forestry pests. Their capacity to adapt rapidly to new environments, evolve resistance to control measures (North et al. 2024), and expand their ranges (Schneider et al. 2022) poses significant challenges for biodiversity conservation, food security, and economic stability. Understanding the ecological and evolutionary processes that govern their adaptation is therefore also essential for developing sustainable management strategies and mitigating future impacts (see, e.g., Chen et al. 2021).

The genus *Ostrinia* (Hübner, 1825; Lepidoptera: Crambidae) has long attracted interest since several of its 23 described species are recognized as major pests of agricultural crops. The European corn borer *O. nubilalis* is among the most devastating pests of maize (*Zea mays* L.) and is responsible for annual yield losses in the United States that exceed $1 billion (Mason et al. 1996). In southeast Asia and Australia, infestations from the Asian corn borer *O. furnacalis* cause heavy damage in several crops, including corn, locally causing up to 30% yield reduction (Zhou et al. 1995). Although a less economically significant pest, the Adzuki bean borer *O. scapulalis* is a phytophagous moth that feeds on several dicotyledonous plants, including hop (*Humulus lupulus*), mugwort (*Artemisia vulgaris*), and hemp (*Cannabis sativa*). *O. scapulalis* is widely distributed across Eurasia and, in Europe, co-occurs with its sibling species *O. nubilalis* (Frolov et al. 2007). Both species show low overall genetic divergence (Malausa et al. 2007) and display patterns of host plant specialization (Bethenod et al. 2005; Calcagno et al. 2007, 2017; Orsucci et al. 2016; Thomas et al. 2003), making the *O. nubilalis*-*O. scapulalis* system a model for the study of divergence in phytophagous insects (Malausa et al. 2005; Streiff et al. 2014).

While multiple chromosome-level genome assemblies have recently been generated for both *O. nubilalis* (Boyes et al. 2025; Ding et al. 2025) and *O. furnacalis* (Peng et al. 2023; Dai et al. 2024; Liu et al. 2025), the only resource available for *O. scapulalis* was based on short read sequencing (Gschloessl et al. 2018). This resulted in a fragmented and incomplete assembly with 50,738 scaffolds, a N50 of ∼29 kbp, and BUSCO scores below 70%, with approximately 60 Mbp of sequence missing compared to the more recent *O. nubilalis* and *O. furnacalis* assemblies. Here, we combined PacBio long read sequencing and Hi-C to produce the first high-quality, chromosome-level genome assembly for *O. scapulalis*, which we annotated using previously released and new RNAseq data, along with protein evidence and *ab initio* predictions. The quality of this resource is on par with other *Ostrinia* chromosome-level assemblies published to date and it will boost research on host plant adaptation and speciation in the *Ostrinia* genus.

## Methods

### Biological materials

Diapausing *O. scapulalis* caterpillars were collected from mugwort stems in February 2023 near Lesquin, France (50°34′ N, 3°6′ E). Individuals were brought to the laboratory to initiate diapause termination at 25°C under a 16:8 light-dark cycle. Because both *O. scapulalis* and *O. nubilalis* occur in the sampling area and cannot be reliably distinguished morphologically, species identification was performed using molecular markers on a subset of 40 individuals (ca. 10% of the total samples). First, samples were genotyped at eight microsatellite loci diagnostic for species delimitation in the system (see Bourguet et al. 2014 and references therein). Second, *O. scapulalis* and *O. nubilalis* produce sex pheromone blends dominated by either E- or Z-Δ11-tetradecenyl acetate (in short, E and Z pherotypes), a polymorphism controlled by the *pgfar* gene, which encodes a pheromone gland–expressed fatty-acyl reductase (Lassance et al. 2010). Individual pherotypes were determined using the PCR-based assay developed by Coates et al. (2013), which relies on three diagnostic single-nucleotide polymorphisms within *pgfar*. Both microsatellite genotyping and pherotyping consistently assigned all individuals to *O. scapulalis*.

Samples were monitored every two days to remove dead or parasitized individuals, and pupae were collected after approximately two weeks. in Lepidoptera, females are the heterogametic sex (ZW), and as we wanted to obtain sufficient read depth for assembling the Z chromosome, two male pupae (ZZ) were retained for downstream steps. We carried sex determination of pupae using multiplex PCR amplification of two microsatellite markers located on the Z and W chromosomes, respectively (Coates and Hellmich 2003). Finally, combined genotyping and pherotyping assays described above were repeated on these two individuals to confirm their *O. scapulalis* assignment.

### Nucleic acid library preparation

For long read sequencing, separate DNA extractions were performed using one-third of a fresh pupa from each of the two individuals. Extractions were carried out with QIAGEN Genomic-tip 20/G kits according to the manufacturer’s instructions. Briefly, we performed a RNase treatment, followed by overnight proteinase K digestion at 54 °C. The DNA pellet was resuspended in 30 µl of Qiagen ultrapure H_2_O, kept overnight at 4 °C, then stored for 48 h at –20 °C before performing quality controls. DNA was quantified using a Qubit fluorometer (Life Technologies), purity was assessed with NanoDrop spectrophotometer, and integrity was evaluated by agarose gel electrophoresis. The highest-quality extraction yielded 6.9 µg of DNA, with an average fragment size of approximately 93 kbp, and was selected for long-read sequencing.

Long read and Hi-C library preparation and sequencing were performed at GeT-PlaGe core facility, INRAE Toulouse, according to the manufacturer’s instructions (“Procedure & Checklist – Preparing whole genome and metagenome libraries using SMRTbell® prep kit 3.0”). At each step, DNA was quantified using the Qubit dsDNA HS

Assay Kit (Life Technologies). DNA purity was tested using the NanoDrop (Thermofisher) and size distribution and degradation assessed using the Femto pulse Genomic DNA 165 kb Kit (Agilent). Purification steps were performed using AMPure PB beads (Pacific Biosciences) and SMRTbell cleanup beads (Pacific Biosciences). Before shearing, we made a repair step with the “SMRTbell Damage Repair Kit SPV3” (Pacific Biosciences), after which 5 μg of DNA were purified and sheared at 20 kbp using the Megaruptor system (Diagenode). Single strand overhangs removal, both DNA and end damage repair and nuclease steps were performed using the SMRTbell® prep kit 3.0. A size selection step at 9 kbp was performed on the BluePippin Size Selection system (Sage Science) following the “0.75%, DF Marker S1 high-pass 6kb-10kb V3” protocol. The first fraction was discarded and the second was recovered in manual mode to obtain a 23 kb library. Using Binding kit 3.2 and sequencing kit 2.0, the library was sequenced by diffusion loading with the adaptive method on a single SMRTcell on Sequel2 instrument at 90 pM with a 2-hour pre-extension and a 30-hour movie.

For Hi-C, a unique library was built from a single pupa from the same population using the Arima-High Coverage Hi-C Kit according to the manufacturer’s instructions. Briefly, we crosslinked and homogenized tissues from a single frozen pupa. Then, chromatin was digested by four restriction enzymes and chromatin ends were biotinylated and ligated by proximity ligation. After reverse crosslink and purification, 200 ng of DNA was used for library preparation. Library sequencing was performed using the Arima Library Prep kit on Illumina NovaSeq 6000, targeting 150 million 150-bp paired-end reads.

### Genome assembly and scaffolding

Unless stated otherwise, all tools were run using default parameters. We computed *k*-mer spectra using KMC v3.2.1 (Kokot et al. 2017) and estimated heterozygosity, genome size, and repetitiveness using GenomeScope v2 (Ranallo-Benavidez et al. 2020). Long reads were assembled using HiFiasm v0.16.1-r375 (Cheng et al. 2021) and completeness was assessed using BUSCO v5.4.4 (Manni et al. 2021) with the Lepidoptera ODB10 dataset (*n* = 5286 orthologs). As the BUSCO duplication rate exceeded 10% (see Results and Discussion), we ran Purge Haplotigs v1.1.2 to flag allelic contigs (Roach et al. 2018) and controlled with BUSCO to ensure the assembly was indeed purged. The mitochondrial genome was extracted from PacBio HiFi reads and annotated using MitoHiFi v3.2 (https://github.com/marcelauliano/MitoHiFi). We used BlobToolKit v4.1.5 (Challis et al. 2020) to ensure no contaminant sequences were present, before assembling contigs into pseudo-chromosomes using the nf-core/hic pipeline v2.1.0-gfe4ac65 (Ewels et al. 2020; Servant et al. 2023). Misjoins were manually corrected using Juicebox v1.11.08 (Durand et al. 2016).

### Gene expression data

We took advantage of two previously published RNA-seq datasets, each consisting of paired-end 100 bp reads generated on Illumina HiSeq 2000 platforms. The first dataset was originally produced to annotate the first *O. scapulalis* assembly (Gschloessl et al. 2018) and comprises seven libraries produced from egg masses, fifth-instar larva whole bodies, fifth-instar larval hemolymph, and adult heads/thoraxes and abdomens (both male and female, NCBI BioProject PRJNA390510). In total, this dataset contains 325 million reads. The second dataset was produced by Orsucci et al. (2018) for a reciprocal transplant experiment, in which *O. scapulalis* individuals were reared on either maize or mugwort. It includes three pools of 20 fourth-instar larvae per treatment, with balanced sex ratios in each pool, totaling 253 million reads (NCBI BioProject PRJNA392376).

Additionally, we produced new RNA-seq data, targeting specifically organs involved in chemoreception. To do so, *O. scapulalis* pupae were produced in the lab by propagating the sampled population on an artificial diet (adapted from Poitout et al. 1972), for several generations between June and November 2023. Around 50 female adults emerging from these pupae were dissected to produce two samples: antennae and gustatory organs (proboscis, palps, first pair of legs, ovipositor). Tissues were collected and immediately placed in TRIzol™ reagent (Thermo Fisher Scientific, Waltham, MA, USA). RNA extraction was conducted following the manufacturer protocol. RNA purity and concentration were measured on a NanoDropTM ND-2000 spectrophotometer (Thermo Fisher Scientific). Library construction and sequencing were carried out at the Novogene company (paired-end 150 bp reads). Briefly, messenger RNAs were purified from total RNAs using poly-T oligo-attached magnetic beads. After fragmentation, the first strand cDNAs were synthesized using random hexamer primers followed by the second strand cDNA synthesis. The libraries were ready after end repair, A-tailing, adapter ligation, size selection, amplification, and purification. The number of read pairs produced were 64.6 million and 69 million for antennae and gustatory organs, respectively.

### Repeat and gene annotation

The next steps were carried on the assembled contigs, i.e., prior to scaffolding with Hi-C data. A *de novo* transposable element (TE) consensus library was built with Repeatmodeler v2.0.3 (Flynn et al. 2020) and was used to mask TE sequences and repeats using Repeatmasker v4.1.2 (Smit et al. 2013). Protein-coding genes in the *O. scapulalis* genome were predicted after soft masking by integrating homology-based, transcript-based, and ab initio approaches with the Braker v3.0.3 pipeline (Gabriel et al. 2024). All protein data available for Arthropoda were downloaded from OrthoDB v11 (https://bioinf.uni-greifswald.de/bioinf/partitioned_odb11, last accessed November 2nd, 2025) and aligned using ProtHint (Brůna et al. 2020). RNA-seq data were trimmed using Fastp 0.22.0 (Chen et al. 2018) and aligned against the hard-masked genome using STAR v2.7.10b (Dobin et al. 2013). Both RNA-seq- and protein-derived hints were used to train GeneMark-ETP, which predictions were in turn used to train Augustus 3.4.0 (Keller et al. 2011). After annotation, UTRs were added to the gene models with GUSHR (https://github.com/Gaius-Augustus/GUSHR). Proteome completeness was evaluated using BUSCO v5.4.4, OMARK v0.3.0 (Nevers et al. 2025) and PSAURON v1.0.6 (Sommer et al. 2025). Finally, automatic functional annotation was carried out from predicted protein sequences with the Blast2GO v6.0 pipeline (Götz et al. 2008).

## Results and Discussion

PacBio sequencing generated 1,956,531 HiFi reads with a mean length of 17.3 kbp, totalling 33.8 Gbp of data. Genome size was estimated at 448,775,962 bp using *k*-mer analysis (*k* = 25), and *k*-mer frequency profiles showed a bimodal distribution, indicating a heterozygous genome, as expected for a field-collected individual (Fig. 1a). The raw assembly produced by HiFiasm spanned 547,829,040 bp across 104 contigs, an N50 of 16.1 Mbp, L50 of 15, a maximum contig length of 28.3 Mbp (Fig. 1b, the longest contig corresponding to the Z chromosome) and a GC content of 37.7%. BUSCO analysis indicated high completeness but notable duplication in the raw assembly (C:98.5%[S:87.6%,D:10.9%],F:0.5%,M:1.0%, *n*=5,286), consistent with the *k*-mer results. Purging haplotigs reduced the total genome size to 485 Mbp in 48 contigs, increased the N50 to 16.5 Mbp and decreased the L50 to 13. A subsequent BUSCO analysis showed that purging removed most duplication while maintaining assembly completeness (C:98.5%[S:97.6%,D:0.9%],F:0.5%,M:1.0%, *n*=5,286). The Hi-C data scaffolded the 48 contigs into the expected 31 pseudo-chromosomes (Guthrie et al. 1965) totaling 475,881,831 bp, with chromosome sizes ranging from 6,362,247 bp to 28,341,256 bp (Fig. 1c, Table 1). A complete circularized mitogenome of 15,247 bp was successfully assembled with MitoHiFi. Overall, the contiguity and completeness of the genome is comparable to the other chromosome-level assemblies released so far for the *Ostrinia* genus (Table 1).

**Table 1.**
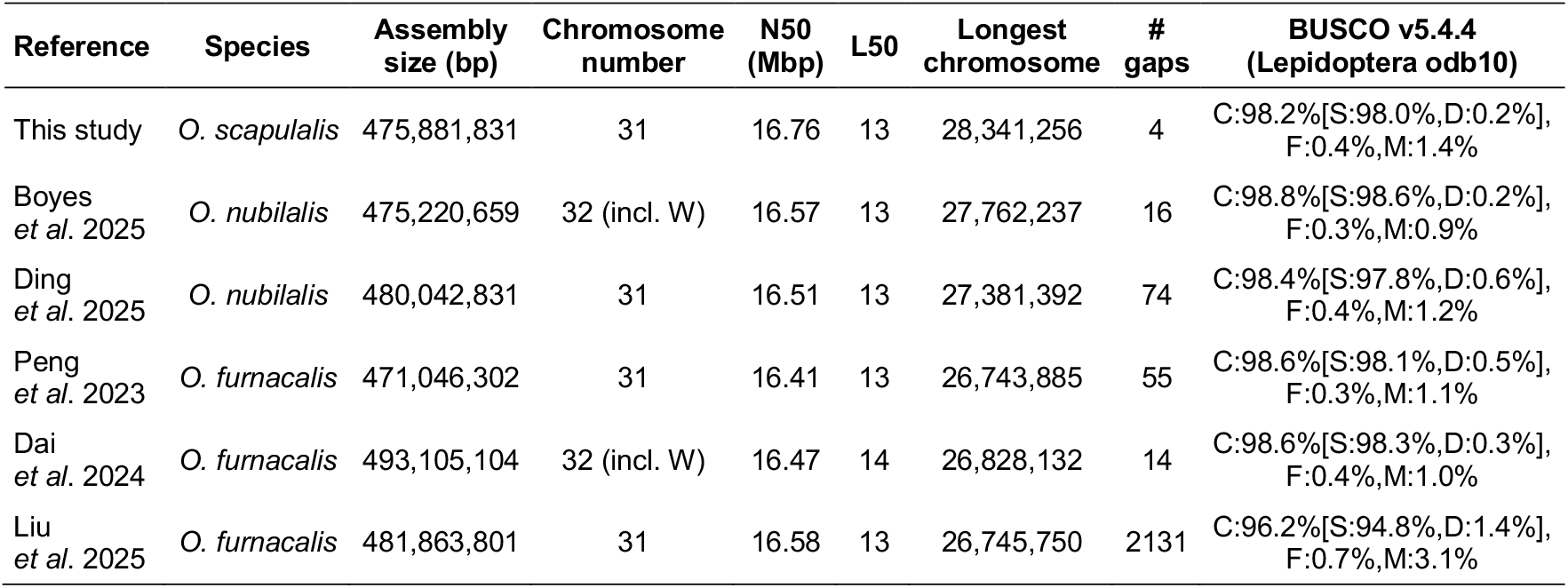
Comparison of the main statistics for *Ostrinia* chromosome-level assemblies published to date.

**Figure 1.**
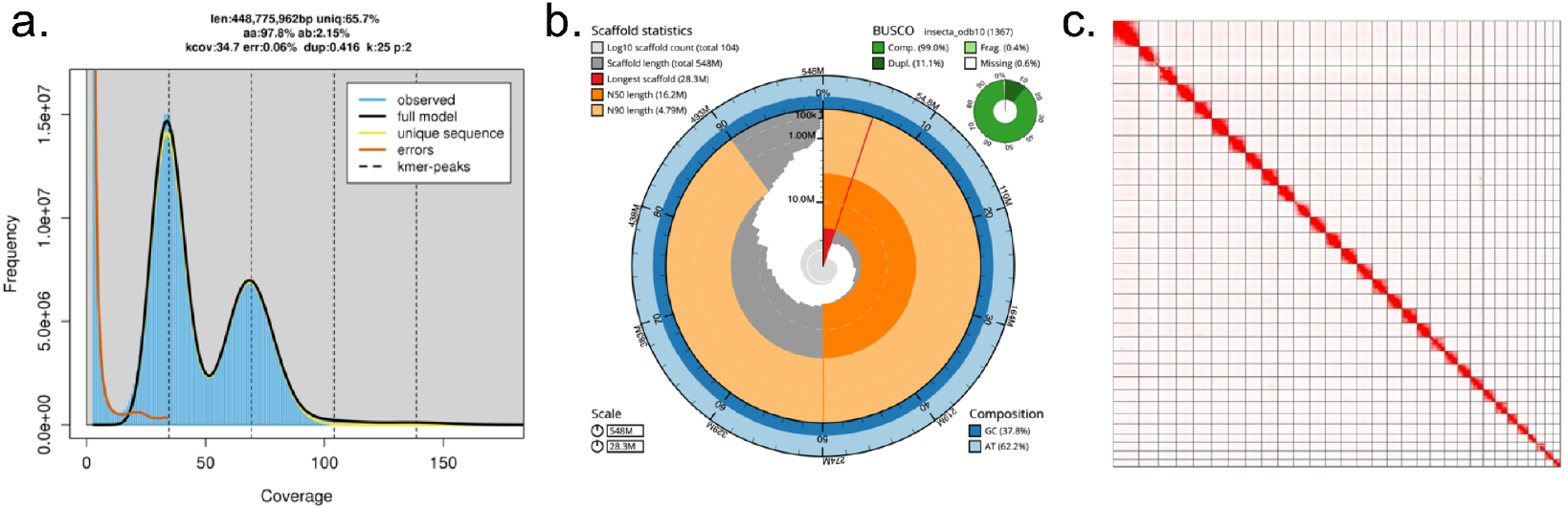
Description of the chromosome-level assembly of the *O. scapulalis* genome. **a**. Distribution of 25-mers generated from the PacBio HiFi data of a single *O. scapulalis* male and visualized with GenomeScope v2. **b**. Snailplot built on contigs using the BlobToolKit v4.1.5 after assembly with HiFiasm v v0.16.1-r375. **c**. Genome-wide Hi-C interaction maps displaying the expected 31 chromosomes of *O. scapulalis*. Interaction frequencies are represented by color intensity, with higher interaction levels (deeper colors) correlating with closer spatial proximity.

After consensus TE library construction, Repeatmasker identified and classified 205,302,855 bp (42.32%) as repeats, with a majority of sequences assigned to Long Interspersed Nuclear Elements (LINEs, 10.39%, Table 2). Our gene annotation pipeline identified 14,261 gene models and 19,888 transcripts (Table 2), with good proteome completeness (BUSCO: C:95.0%[S:93.6%,D:1.4%],F:0.6%,M:4.4%, *n*=5,286; OMARK completeness: 94.4%, PSAURON score: 97.9). Automatic functional annotation carried with BLAST2GO assigned functional labels to 15,795 transcripts (79.42%), with only 335 transcripts (1.7%) without any significant BLAST hit.

**Table 2.** Main annotation statistics for the chromosome-level assembly of *O. scapulalis*.

|  |  |  |
| --- | --- | --- |
| **Gene annotation** | Total number of gene models | 14,152 |
|  | Mean gene length (bp) | 32,086 |
|  | Average number of exons per gene | 8.53 |
|  | Cumulative exon length (Mbp, %) | 60.90 (12.8) |
|  | Number of isoforms | 19,888 |
|  | Average number of isoforms per gene | 1.40 |
|  | Mean protein length (amino acids) | 536 |
| **Repeat annotation** | Percentage of genome masked | 42.32 |
|  | Retroelements (%) | 14.27 |
|  | LINEs (%) | 10.39 |
|  | LTR elements (%) | 3.28 |
|  | SINEs (%) | 0.60 |
|  | DNA transposons (%) | 3.61 |
|  | Unclassified (%) | 17.70 |

## Conclusion

Here, we present a high-quality, annotated genome assembly for *O. scapulalis*. This new resource contributes to a robust foundation for investigating the genomic mechanisms underlying host plant adaptation in *Ostrinia*, a genus that includes two of the most destructive maize pests worldwide. By offering a substantial improvement over previously available genomic data for *O. scapulalis*, this assembly will facilitate comparative analyses across *Ostrinia* species, enabling researchers to dissect the genetic basis of ecological specialization, host shifts, and life-history diversification. Moreover, the availability of this high-quality genome will help clarify the evolutionary history of divergence between *O. scapulalis* and its sibling species *O. nubilalis*, a long-standing question in evolutionary biology (Bierne et al. 2011; Bourguet et al. 2014; Kunerth et al. 2022). This resource will support future studies aimed at understanding how genomic architecture, gene flow, and selection interact during the early stages of speciation.

## Acknowledgements

This work was performed in collaboration with the GeT core facility, Toulouse, France (GeT, https://doi.org/10.15454/1.5572370921303193E12). GeT core facility was supported by France Génomique National infrastructure, funded as part of “Investissement d’avenir” program managed by Agence Nationale pour la Recherche (contract ANR-10-INBS-09). We acknowledge the genotoul bioinformatics platform Toulouse Occitanie (Bioinfo Genotoul, https://doi.org/10.15454/1.5572369328961167E12) for providing computing and storage resources.

## Data Availability

Contig sequences and the chromosome-level assembly were deposited in the European Nucleotide Archive (###). The assembly and its annotation are available through the BioInformatics Platform for Agroecosystem Arthropods (bipaa.genouest.org).

## Literature cited

Bethenod, M-T, Y. Thomas, F. Rousset, et al. 2005. “Genetic Isolation between Two Sympatric Host Plant Races of the European Corn Borer, Ostrinia Nubilalis Hübner. II: Assortative Mating and Host-Plant Preferences for Oviposition.” Heredity 94 (2): 264–270.

Bierne, Nicolas, John Welch, Etienne Loire, François Bonhomme, and Patrice David. 2011. “The Coupling Hypothesis: Why Genome Scans May Fail to Map Local Adaptation Genes.” Molecular Ecology 20 (10): 2044–2072.

Bourguet, Denis, Sergine Ponsard, Rejane Streiff, et al. 2014. “‘Becoming a Species by Becoming a Pest’ or How Two Maize Pests of the Genus Ostrinia Possibly Evolved through Parallel Ecological Speciation Events.” Molecular Ecology 23 (2): 325–342.

Boyes, Douglas, David C. Lees, Brad S. Coates, et al. 2025. “The Genome Sequence of the European Corn Borer, Ostrinia nubilalis Hübner, 1796.” Wellcome Open Research 10 (January): 12.

Breeschoten, Thijmen, Corné F. H. van der Linden, Vera I. D. Ros, M. Eric Schranz, and Sabrina Simon. 2022. “Expanding the Menu: Are Polyphagy and Gene Family Expansions Linked across Lepidoptera?” Genome Biology and Evolution 14 (1): evab283.

Brůna, Tomáš, Alexandre Lomsadze, and Mark Borodovsky. 2020. “GeneMark-EP+: Eukaryotic Gene Prediction with Self-Training in the Space of Genes and Proteins.” NAR Genomics and Bioinformatics 2 (2): qaa026.

Calcagno, Vincent, Clémentine Mitoyen, Philippe Audiot, et al. 2017. “Parallel Evolution of Behaviour during Independent Host-Shifts Following Maize Introduction into Asia and Europe.” Evolutionary Applications 10 (9): 881–889.

Calcagno, V., Y. Thomas, and D. Bourguet. 2007. “Sympatric Host Races of the European Corn Borer: Adaptation to Host Plants and Hybrid Performance.” Journal of Evolutionary Biology 20 (5): 1720– 1729.

Challis, Richard, Edward Richards, Jeena Rajan, Guy Cochrane, and Mark Blaxter. 2020. “BlobToolKit - Interactive Quality Assessment of Genome Assemblies.” G3 (Bethesda, Md.) 10 (4): 1361–1374.

Cheng, Haoyu, Gregory T. Concepcion, Xiaowen Feng, Haowen Zhang, and Heng Li. 2021. “Haplotype-Resolved de Novo Assembly Using Phased Assembly Graphs with Hifiasm.” Nature Methods 18 (2): 170–175.

Chen, Shifu, Yanqing Zhou, Yaru Chen, and Jia Gu. 2018. “Fastp: An Ultra-Fast All-in-One FASTQ Preprocessor.” Bioinformatics 34 (17): i884–i890.

Chen, Yanting, Zhaoxia Liu, Jacques Régnière, et al. 2021. “Large-Scale Genome-Wide Study Reveals Climate Adaptive Variability in a Cosmopolitan Pest.” Nature Communications 12 (1): 7206.

Coates, Brad S., and Richard L. Hellmich. 2003. “Two Sex-Chromosome-Linked Microsatellite Loci Show Geographic Variance among North American Ostrinia Nubilalis.” Journal of Insect Science 3 (1): 29.

Coates, Brad S., Holly Johnson, Kyung-Seok Kim, et al. 2013. “Frequency of Hybridization between Ostrinia Nubilalis E-and Z-Pheromone Races in Regions of Sympatry within the United States.” Ecology and Evolution 3 (8): 2459–2470.

Dai, Wenting, Judith E. Mank, and Liping Ban. 2024. “Gene Gain and Loss from the Asian Corn Borer W Chromosome.” BMC Biology 22 (1): 102.

Ding, Xinhua, Yue Zhang, Xiaowu Wang, et al. 2025. “Chromosome-Level Genome Assembly of Z Strain European Corn Borer Ostrinia Nubilalis (Lepidoptera: Crambidae).” Scientific Data 12 (1): 365.

Dobin, Alexander, Carrie A. Davis, Felix Schlesinger, et al. 2013. “STAR: Ultrafast Universal RNA-Seq Aligner.” Bioinformatics 29 (1): 15–21.

Drès, Michele, and James Mallet. 2002. “Host Races in Plant-Feeding Insects and Their Importance in Sympatric Speciation.” Philosophical Transactions of the Royal Society of London. Series B, Biological Sciences 357 (1420): 471–492.

Durand, Neva C., James T. Robinson, Muhammad S. Shamim, et al. 2016. “Juicebox Provides a Visualization System for Hi-C Contact Maps with Unlimited Zoom.” Cell Systems 3 (1): 99–101.

Ewels, Philip A., Alexander Peltzer, Sven Fillinger, et al. 2020. “The Nf-Core Framework for Community-Curated Bioinformatics Pipelines.” Nature Biotechnology 38 (3): 276–278.

Flynn, Jullien M., Robert Hubley, Clément Goubert, et al. 2020. “RepeatModeler2 for Automated Genomic Discovery of Transposable Element Families.” Proceedings of the National Academy of Sciences of the United States of America 117 (17): 9451–9457.

Forbes, Andrew A., Sara N. Devine, Alaine C. Hippee, et al. 2017. “Revisiting the Particular Role of Host Shifts in Initiating Insect Speciation.” Evolution; International Journal of Organic Evolution 71 (5): 1126–1137.

Forister, M. L., L. A. Dyer, M. S. Singer, J. O. Stireman 3rd, and J. T. Lill. 2012. “Revisiting the Evolution of Ecological Specialization, with Emphasis on Insect-Plant Interactions.” Ecology 93 (5): 981–991.

Frolov, Andrei N., Denis Bourguet, and Sergine Ponsard. 2007. “Reconsidering the Taxonomy of Several Ostrinia Species in the Light of Reproductive Isolation: A Tale for Ernst Mayr.” Biological Journal of the Linnean Society. Linnean Society of London 91 (1): 49–72.

Futuyma, D., and Gabriel Moreno. 1988. “The Evolution of Ecological Specialization.” Annual Review of Ecology, Evolution, and Systematics 19: 207–233.

Futuyma, Douglas J., and Anurag A. Agrawal. 2009. “Macroevolution and the Biological Diversity of Plants and Herbivores.” Proceedings of the National Academy of Sciences of the United States of America 106 (43): 18054–18061.

Gabriel, Lars, Tomáš Brůna, Katharina J. Hoff, et al. 2024. “BRAKER3: Fully Automated Genome Annotation Using RNA-Seq and Protein Evidence with GeneMark-ETP, AUGUSTUS, and TSEBRA.” Genome Research 34 (5): 769–777.

Götz, Stefan, Juan Miguel García-Gómez, Javier Terol, et al. 2008. “High-Throughput Functional Annotation and Data Mining with the Blast2GO Suite.” Nucleic Acids Research 36 (10): 3420–3435.

Gschloessl, B., F. Dorkeld, P. Audiot, A. Bretaudeau, C. Kerdelhué, and R. Streiff. 2018. “De Novo Genome and Transcriptome Resources of the Adzuki Bean Borer Ostrinia Scapulalis (Lepidoptera: Crambidae).” Data in Brief 17 (April): 781–787.

Guthrie, W. D., E. J. Dollinger, and J. F. Stetson. 1965. “Chromosome Studies of the European Corn Borer, Smartweed Borer, and Lotus Borer (Pyralidae)1.” Annals of the Entomological Society of America 58 (1): 100–105.

Jaenike, John. 1990. “Host Specialization in Phytophagous Insects.” Annual Review of Ecology and Systematics 21 (1): 243–273.

Jousselin, Emmanuelle, and Marianne Elias. 2019. “Testing Host-Plant Driven Speciation in Phytophagous Insects: A Phylogenetic Perspective.” In arXiv [q-bio.PE]. February 25. arXiv. 10.20944/preprints201902.0215.v1.

Keller, Oliver, Martin Kollmar, Mario Stanke, and Stephan Waack. 2011. “A Novel Hybrid Gene Prediction Method Employing Protein Multiple Sequence Alignments.” Bioinformatics (Oxford, England) 27 (6): 757–763.

Kokot, Marek, Maciej Dlugosz, and Sebastian Deorowicz. 2017. “KMC 3: Counting and Manipulating K-Mer Statistics.” Bioinformatics (Oxford, England) 33 (17): 2759–2761.

Kunerth, Henry D., Steven M. Bogdanowicz, Jeremy B. Searle, et al. 2022. “Consequences of Coupled Barriers to Gene Flow for the Build-up of Genomic Differentiation.” Evolution; International Journal of Organic Evolution 76 (5): 985–1002.

Lassance, Jean-Marc, Astrid T. Groot, Marjorie A. Liénard, et al. 2010. “Allelic Variation in a Fatty-Acyl Reductase Gene Causes Divergence in Moth Sex Pheromones.” Nature 466 (7305): 486–489.

Liu, Kaiqiang, Tiantao Zhang, Yongjun Zhang, Zhenying Wang, and Kanglai He. 2025. “Chromosome-Level Genome of a Multivoltine Biotype Ostrinia Furnacalis Strain.” Scientific Data 12 (1): 881.

Malausa, T., A. Dalecky, S. Ponsard, et al. 2007. “Genetic Structure and Gene Flow in French Populations of Two Ostrinia Taxa: Host Races or Sibling Species?” Molecular Ecology 16 (20): 4210–4222.

Malausa, Thibaut, Marie-Thérèse Bethenod, Arnaud Bontemps, Denis Bourguet, Jean-Marie Cornuet, and Sergine Ponsard. 2005. “Assortative Mating in Sympatric Host Races of the European Corn Borer.” Science (New York, N.Y.) 308 (5719): 258–260.

Manni, Mosè, Matthew R. Berkeley, Mathieu Seppey, Felipe A. Simão, and Evgeny M. Zdobnov. 2021. “BUSCO Update: Novel and Streamlined Workflows along with Broader and Deeper Phylogenetic Coverage for Scoring of Eukaryotic, Prokaryotic, and Viral Genomes.” Molecular Biology and Evolution 38 (10): 4647–4654.

Mason, C. E., M. E. Rice, and D. D. Calvin, eds. 1996. European Corn Borer Ecology and Management. Iowa State University.

McCulloch, Graham A., and Jonathan M. Waters. 2023. “Rapid Adaptation in a Fast-Changing World: Emerging Insights from Insect Genomics.” Global Change Biology 29 (4): 943–954.

Nevers, Yannis, Alex Warwick Vesztrocy, Victor Rossier, et al. 2025. “Quality Assessment of Gene Repertoire Annotations with OMArk.” Nature Biotechnology 43 (1): 124–133.

North, Henry L., Zhen Fu, Richard Metz, et al. 2024. “Rapid Adaptation and Interspecific Introgression in the North American Crop Pest Helicoverpa Zea.” Molecular Biology and Evolution 41 (7). 10.1093/molbev/msae129.

Orsucci, M., P. Audiot, F. Dorkeld, et al. 2018. “Larval Transcriptomic Response to Host Plants in Two Related Phytophagous Lepidopteran Species: Implications for Host Specialization and Species Divergence.” BMC Genomics 19 (1). 10.1186/s12864-018-4589-x.

Orsucci, M., P. Audiot, A. Pommier, et al. 2016. “Host Specialization Involving Attraction, Avoidance and Performance, in Two Phytophagous Moth Species.” Journal of Evolutionary Biology 29 (1): 114–125.

Peng, Yan, Minghui Jin, Zhimin Li, et al. 2023. “Population Genomics Provide Insights into the Evolution and Adaptation of the Asia Corn Borer.” Molecular Biology and Evolution 40 (5). 10.1093/molbev/msad112.

Poitout, S., R. Bues, and C. L. E. Rumeur. 1972. “Élevage Sur Milieu Artificiel Simple de Deux Noctuelles Parasites Du Coton Earias Insulana et Spodoptera Littoralis.” Entomologia Experimentalis et Applicata 15 (3): 341–350.

Ranallo-Benavidez, T. Rhyker, Kamil S. Jaron, and Michael C. Schatz. 2020. “GenomeScope 2.0 and Smudgeplot for Reference-Free Profiling of Polyploid Genomes.” Nature Communications 11 (1): 1432.

Roach, Michael J., Simon A. Schmidt, and Anthony R. Borneman. 2018. “Purge Haplotigs: Allelic Contig Reassignment for Third-Gen Diploid Genome Assemblies.” BMC Bioinformatics 19 (1): 460.

Schneider, Léonard, Martine Rebetez, and Sergio Rasmann. 2022. “The Effect of Climate Change on Invasive Crop Pests across Biomes.” Current Opinion in Insect Science 50 (100895): 100895.

Servant, Nicolas, Nf-Core Bot, Phil Ewels, et al. 2023. Nf-Core/hic: Nf-Core/hic v2.1.0. Zenodo. 10.5281/ZENODO.7994878.

Simon, Jean-Christophe, Emmanuelle d’Alençon, Endrick Guy, et al. 2015. “Genomics of Adaptation to Host-Plants in Herbivorous Insects.” Briefings in Functional Genomics 14 (6): 413–423.

Smit, A. F., R. Hubley, and P. Green. 2013. “Repeat-Masker Open-4.0.” http://www.repeatmasker.org.

Sommer, Markus J., Aleksey V. Zimin, and Steven L. Salzberg. 2025. “PSAURON: A Tool for Assessing Protein Annotation across a Broad Range of Species.” NAR Genomics and Bioinformatics 7 (1): qae189.

Streiff, Réjane, Brigitte Courtois, Serge Meusnier, and Denis Bourguet. 2014. “Genetic Mapping of Two Components of Reproductive Isolation between Two Sibling Species of Moths, Ostrinia Nubilalis and O. Scapulalis.” Heredity 112 (4): 370–381.

Thomas, Yan, Marie-Thérèse Bethenod, Laurent Pelozuelo, Brigitte Frérot, and Denis Bourguet. 2003. “Genetic Isolation between Two Sympatric Host-Plant Races of the European Corn Borer, Ostrinia Nubilalis Hübner. I. Sex Pheromone, Moth Emergence Timing, and Parasitism.” Evolution; International Journal of Organic Evolution 57 (2): 261–273.

Zhou, D. R., K. L. He, Wang Z. Y., et al., eds. 1995. Asian Corn Borer and Its Integrated Management. Golden Shield Press.

